# CellMAPS: A no-code model-based customisable multiplex image analysis workflow

**DOI:** 10.1101/2025.10.07.680848

**Authors:** Éanna Fennell, Colm Brandon, Steve Boßelmann, Stephen E. Ryan, Amandeep Singh, Aoife Hennessy, Aisling M. Ross, Ciara I. Leahy, Matthew R. Pugh, Nadezhda Nikulina, Oliver Braubach, Gerald Niedobitek, Graham S. Taylor, Tiziana Margaria, Paul G. Murray

## Abstract

Although multiplex imaging allows simultaneous mapping of complex tissue architectures and cellular phenotypes, the dearth of user-friendly robust image analysis workflows remains a significant limitation to its widespread use, restricting multiplex imaging to laboratories with image analysis expertise. Here, we introduce a no-code model-based customisable suite, CellMAPS, which allows both image processing and spatial analyses by non-specialists. We also developed new tools for tissue de-arraying, image normalisation and cell segmentation to increase usability and reduce analysis subjectivity. We have also introduce new spatial analysis tools for the partition of tissue architectures and cellular microenvironments. We demonstrate the capabilities of CellMAPS in various diseases captured using different multiplex imaging platforms. Overall, CellMAPS provides a platform for inexperienced laboratories to implement multiplex imaging and complex spatial analyses, which ultimately should lead to the broader adoption of these technologies in the biomedical field.

## INTRODUCTION

High-dimensional multiplex imaging has the potential to revolutionise our understanding of tissue disorders. For example, in cancer these technologies allow researchers to map the complex interplay between tumour cells and their immune microenvironment, and to study how these interactions influence cellular phenotype and signalling patterns, and impact on patient response to therapy and clinical outcomes^1–3^. However, the widespread application of these emerging technologies is limited due in part to the difficulties associated with analysis of the images that are generated. These images can span more than 100 channels, capture millions of spatially resolved single cells, and hold over 100GB of data^4^. The scale of this data lends itself towards the generation of new AI-driven analysis techniques to obtain actionable insights from the experimental data. However, bespoke workflows to process and analyse these images are lacking. This is especially true for wet-laboratory researchers who may have little experience in image and bioinformatic analyses. Moreover, research groups must be specifically skilled in multiplex image analysis and be able to install, use and combine various analysis packages. Hence, there is a need for more user-friendly open-source customisable analysis pipelines.

Key to improving access to more user-friendly image analysis will be the development of more generalisable workflows, applicable to diverse types of experiments and tissue types while at the same time reducing the reliance on previous expertise in multiplex image analysis. To achieve this, reducing image variation due to differences in tissue fixation and staining protocols, needs to be addressed. Furthermore, workflows will need to be implemented in environments that are easy to access, utilise and scale with minimal input from the user. There is also a requirement for the further development of spatial concepts and tools to give a more robust tissue characterisation^5–9^. Key amongst these is the quantification of meaningful cellular interactions within tissues on length scales larger than direct interactions.

To tackle these limitations, we present a no-code model-based customisable multiplex imaging analysis suite, CellMAPS (Single-**Cell M**odel-based **A**nalysis **P**ipeline for **S**patial-omics); comprising of HIPPo (**H**igh-plex **I**mmunofluorescence **P**re-processing **P**ipeline **o**rchestrator) for pre-processing and MISSILe (**M**ultiplex **I**mmunofluore**S**cence **S**patial **I**nteraction **L**ibrary) for spatial and phenotypic analyses. HIPPo presents two new packages for user-tuneable tissue de-arraying and image normalisation to generate robust high quality single-cell data for downstream spatial analyses, both enabling increased biological insights over existing tools. MISSILe introduces two new spatial analysis tools for tissue architecture, interactions and immediate microenvironment analysis. CellMAPS was implemented in a bespoke no-code environment, providing an easy-to-use graphical user interface, which can be used off-the-shelf for new users or further developed by experienced bioinformaticians^10^. To show the improved biological insight and ease-of-use capabilities of CellMAPS, we have applied the pipeline to new and existing cancer and COVID-19 image datasets.

## RESULTS

### CellMAPS implements HIPPo & MISSILe in a bespoke no-code environment

To generate a user-friendly, customisable multiplex image analysis workflow, we developed CellMAPS which implements two new modules, HIPPo and MISSILe (Fig. 1). HIPPo, a pre-processing module, generates numerical cellular data from multiplex images captured on any spatial-omics platform through de-arraying, normalisation, cell segmentation and feature extraction. In HIPPo we have developed three new python-based packages, SegArray (for tissue de-arraying), ACE (**A**utomatic **C**ontrast **E**qualisation for normalisation) and Xtractit (for expression and morphological cellular feature extraction). We also combined and integrated previously published cell segmentation models (e.g. DeepCell, CellSeg and CellPose), allowing users to fine tune their multiplex image processing^11–14^. For spatial analysis, we describe MISSILe implemented in R which includes cell filtering and quality control, single-cell annotation (PhenoGraph and CELESTA)^15, 16^, quantification of cell abundances and functional states as well as spatial visualisations (Fig. 1). In addition to making use of previously published spatial concepts (e.g., cellular neighbourhoods)^17^, we have developed Local Aggregation For Tissue Architecture (LAFTA) for tissue architecture analysis. We also present a tool to comprehensively detail the immediate microenvironment of cells.

**Figure 1.**
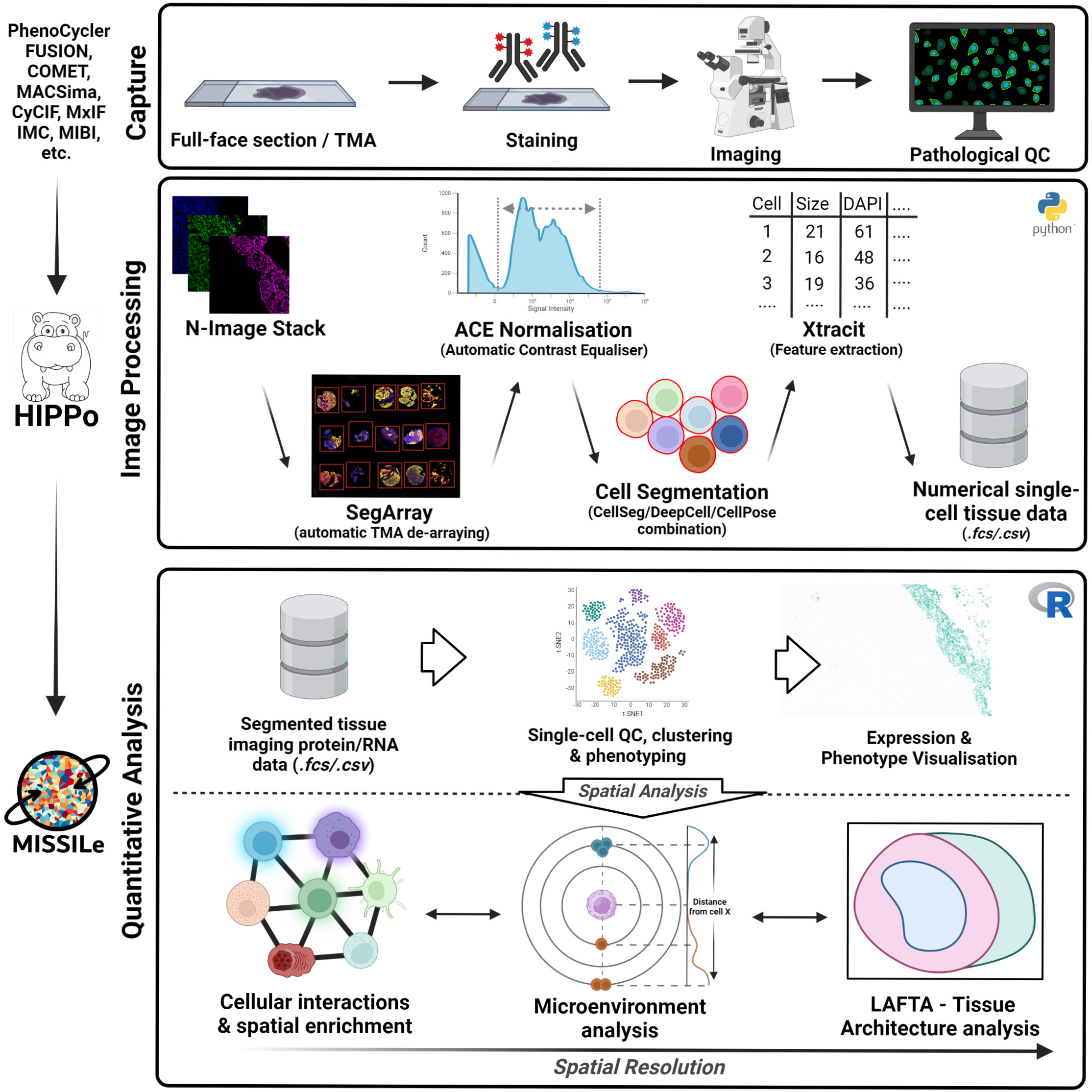
Overview of CellMAPS pipeline. Data captured on any commercially available spatial proteomics platform is passed through the pre-processing module, HIPPo, for de-arraying (if applicable), contrast equalisation, cellular segmentation, data extraction and spatial spill overcompensation. Segmented single-cell data is subsequently quality controlled, clustered, annotated and analysed spatially across various length scales from immediate cellular interactions to local microenvironment to overall tissue architecture (with LAFTA) in MISSILe.

The CellMAPS application consists of a graphical Integrated Modelling Environment (IME), an execution environment and an execution frontend for human-in-the-loop computational processes (Extended Data Fig. 1a). The set of computational processing components (service independent building-blocks; SIBs) are developed using model-driven engineering and micro-service principals, resulting in a pipeline which can automatically decide upon parameters and modules optimal for the input data (Extended Data Fig. 1a & b). The application stack is implemented using container virtualisation, easily deployable both locally and in the cloud and hence is not restricted by the need for large computing infrastructure. Image data is also stored locally or in the cloud. The IME provides pre-made pipeline architectures and enables users to edit existing pipelines.

### SegArray de-arrays irregularly shaped TMAs

Tissue Microarrays (TMAs), which sample areas of multiple tissues onto a single slide, are often used in multiplex imaging^18^. However, TMAs require a de-arraying procedure to split individual cores so that each is analysed individually. We have developed an automatic tissue de-array tool, SegArray, trained on 14 down sampled and manually annotated TMAs. TMAs consisting of lung, placental, lymphoid, breast and nasopharyngeal tissues imaged on the multiple platforms were used for training. Data augmentation techniques generated over one thousand further annotated TMA images which were used to further refine SegArray (see Methods section). All annotated TMAs were used to train a UNET model, resulting in consistent identification of tissue cores in TMAs across a second set of lymphoid and epithelial cancers (Fig. 2a)^19^. However, some tissue types and core sizes, such as 4mm lung cores, did not perform as well as other TMAs. For this we implemented a variable user-selected confidence value which can be optimised to the tissue type and TMA layout as required (Extended Data Fig. 2a). Furthermore, for cases which cannot be handled by SegArray or are predicted incorrectly, manual de-arraying can be employed by either adjusting segmentation predicted masks or drawing new masks with a custom mask annotating tool within CellMAPS (Extended Data Fig. 2b). To test the performance of SegArray, we segmented four TMAs consisting of tissue types not used for training including colorectal cancer (CRC), infectious mononucleosis (IM) tonsil, Hodgkin lymphoma (HL) and Diffuse Large B Cell Lymphoma (DLBCL) with SegArray, Coreograph and QuPath. Given that the primary output of interest is the identification of cores for downstream analysis, we limited the analysis to model predictions, without the use of manual adjustment. However, the ground truth number of cores were defined manually. SegArray correctly identified 93% of cores correctly compared to Coreograph with 73% and QuPath with 70% (Fig. 2b). Furthermore, SegArray had the highest F1 score compared to Coreograph and QuPath (Fig. 2c). SegArray outperformed the other tools considerably on packed TMAs whereas the performance was similar in TMAs with spaced cores. Furthermore, SegArray successfully de-arrayed 34 further tissue types across two unseen TMAs (Extended Data Fig. 2c)^20^. The combination of accurate core identification and manual adjustment results SegArray ensures minimal loss of TMA cores to de-arraying errors, resulting in more samples and cells to extract biological insights from.

**Figure 2.**
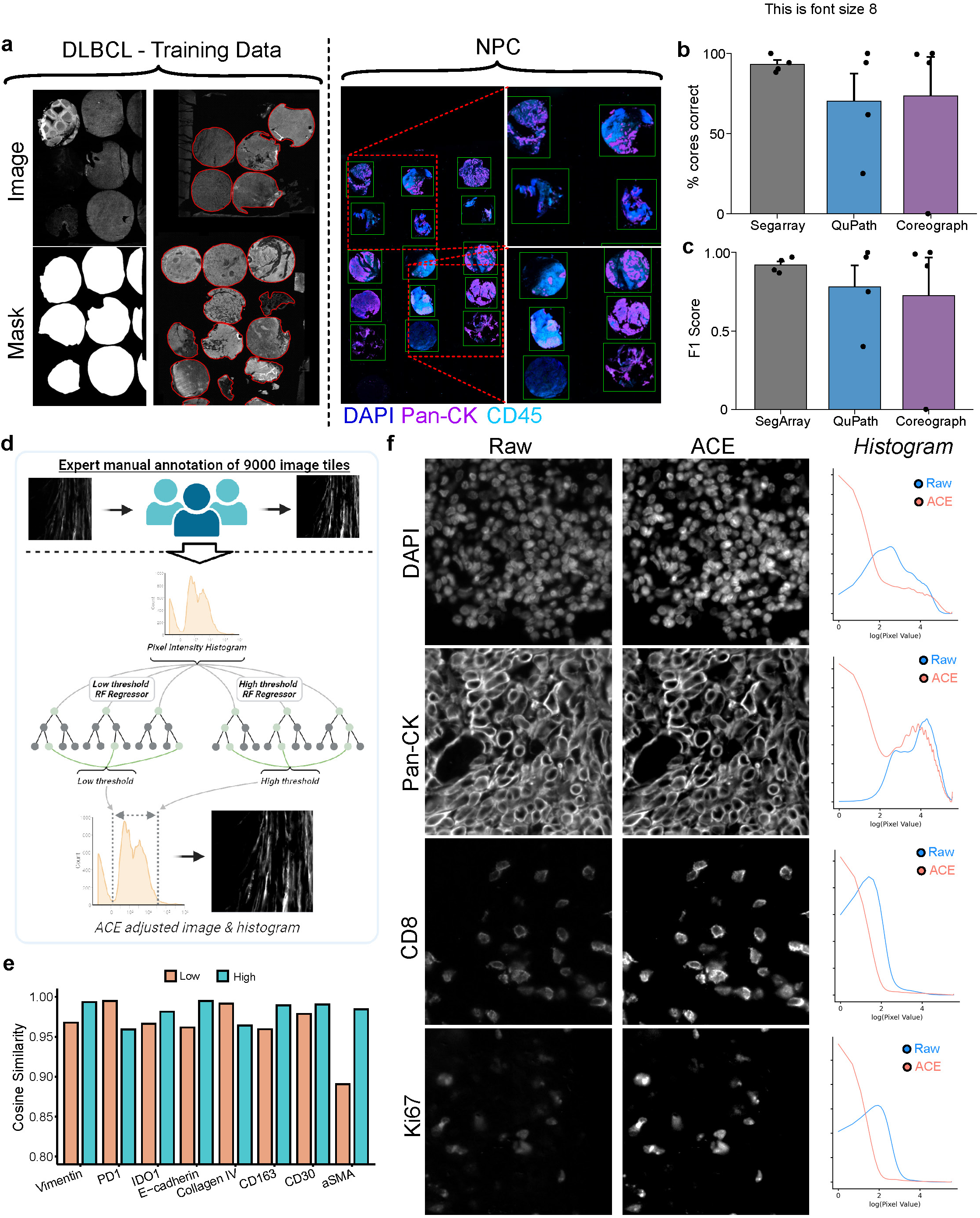
SegArray & ACE model development & benchmarking. a. SegArray is trained on irregular TMA configurations. Segmentation masks were manually generated for training and used to detect cores in lymphoid and epithelial cancers. b-c. Percentage of cores correctly predicted and F1 score in 4 TMAs consisting of various tissue types. Metrics were compared between SegArray, QuPath and Coreograph. d. Automatic Contrast Equaliser (ACE) model schematic showing manually adjusted image data set (n=9000) and the multi-target decision tree strategy to identify lower and upper pixel values for equalisation. e. Image tile input and ACE outputs showing enhanced contrast for low to highly expressed proteins. Pixel expression histograms show larger pixel numbers at low (0) and high (255) values, resulting in a more stretched histogram. f. Cosine similarity of ACE predicted thresholds and expert manual annotations for various structural, stromal and lymphoid proteins.

### ACE increases signal-to-noise ratio

Clinical tissue samples, traditionally formalin-fixed paraffin-embedded (FFPE) processed, show large inter-sample signal heterogeneity, which can pose challenges when attempting to compare protein expression levels between samples. Furthermore, technical staining variation compounds signal heterogeneity across slides (Extended Data Fig. 2c). Numerical integration methods (as used regularly in single-cell RNA sequencing) are difficult to implement in multiplex imaging data due to limited numbers of features. Hence, we developed ACE to identify the optimal signal-to-noise ratio (SNR) from the native images across many protein expression types. ACE consists of a multi-target regression model trained on 9000 manually adjusted image tiles, identifying upper and lower threshold values (Fig. 2b). The image tiles consisted of proteins which consistently have high signal (e.g., pan-cytokeratin), mid signal (e.g., CD20) and low signal (e.g., LAG3) to capture the heterogeneity of expression profiles. Applying ACE to previously unseen image tiles across the three expression profiles showed that ACE increases the contrast at variable amounts depending on the input signal. Although DAPI and pan-cytokeratin channels qualitatively showed little signal-to-noise ratio improvement, CD8 and Ki67 showed a large signal-to-noise improvement. The signal histograms show less noise (negative signal greater than a zero-pixel value) in the ACE images (Fig. 2c). To test the accuracy of the ACE model, we manually annotated a further 320 image tiles across eight proteins and two tissue types and compared the low and high contrast threshold values to the ACE predictions by cosine similarity. Overall, all similarities were calculated at over 0.96 (out of 1) apart from the lower threshold of aSMA (0.89). To utilise ACE at a tissue level, the model was extended to whole tissue images with a computational time of ∼12 seconds per image containing 100M pixels (Extended Data Fig. 2d).

Next, to test if ACE reduces technical variation between mIF images, we stained two immediately adjacent slides from lymph node tissue on the PhenoCycler FUSION with different conditions (pH; 6 & 9, antibody incubation time; 18hr & 24hr, time from staining to imaging post fixation; immediate & 24 hours). We found that the induced technical variation resulted in differences in contrast-to-noise-ratio (CNR) values (taken from the same tissue regions) across multiple channels (Extended Data Fig. 2e). However, applying ACE to both slides reduced the differences in CNR across all channels (Extended Data Fig. 2e).

### ACE normalises signal intensities across heterogeneous mIF datasets

To benchmark ACE across a dataset, we imaged a TMA of 15 nasopharyngeal carcinoma (NPC) samples on the PhenoCycler FUSION with a 42-plex antibody panel. The TMA was de-arrayed with SegArray and subsequently each core was run through ACE. To examine the limits of ACE, we concentrated on low contrast images CD4 and CD38. Plotting the cellular expression (post cell segmentation) across each tissue core, we found ACE increased the contrast between positive and negative expression (Fig. 3a & Extended Data Fig. 3a). The 99^th^ quantile values of ACE increased by 42 and 74 for CD4 and CD38 respectively compared to raw expression with little change in the mean values (5 and 4 for CD4 and CD38 respectively). To test the normalisation of ACE, we calculated the similarity (using 1 – Kolmogorov–Smirnov score) of outlier cores (region 11 for CD4 and 13 for CD38) with all other cores for raw and ACE processing. We found that ACE had higher similarity of outlier cores to other cores for both CD4 and CD38 (Fig. 3b). We also show a statistically significant increase in SNR for both CD4 and CD38 (3 and 40 respectively Fig. 3c).

**Figure 3.**
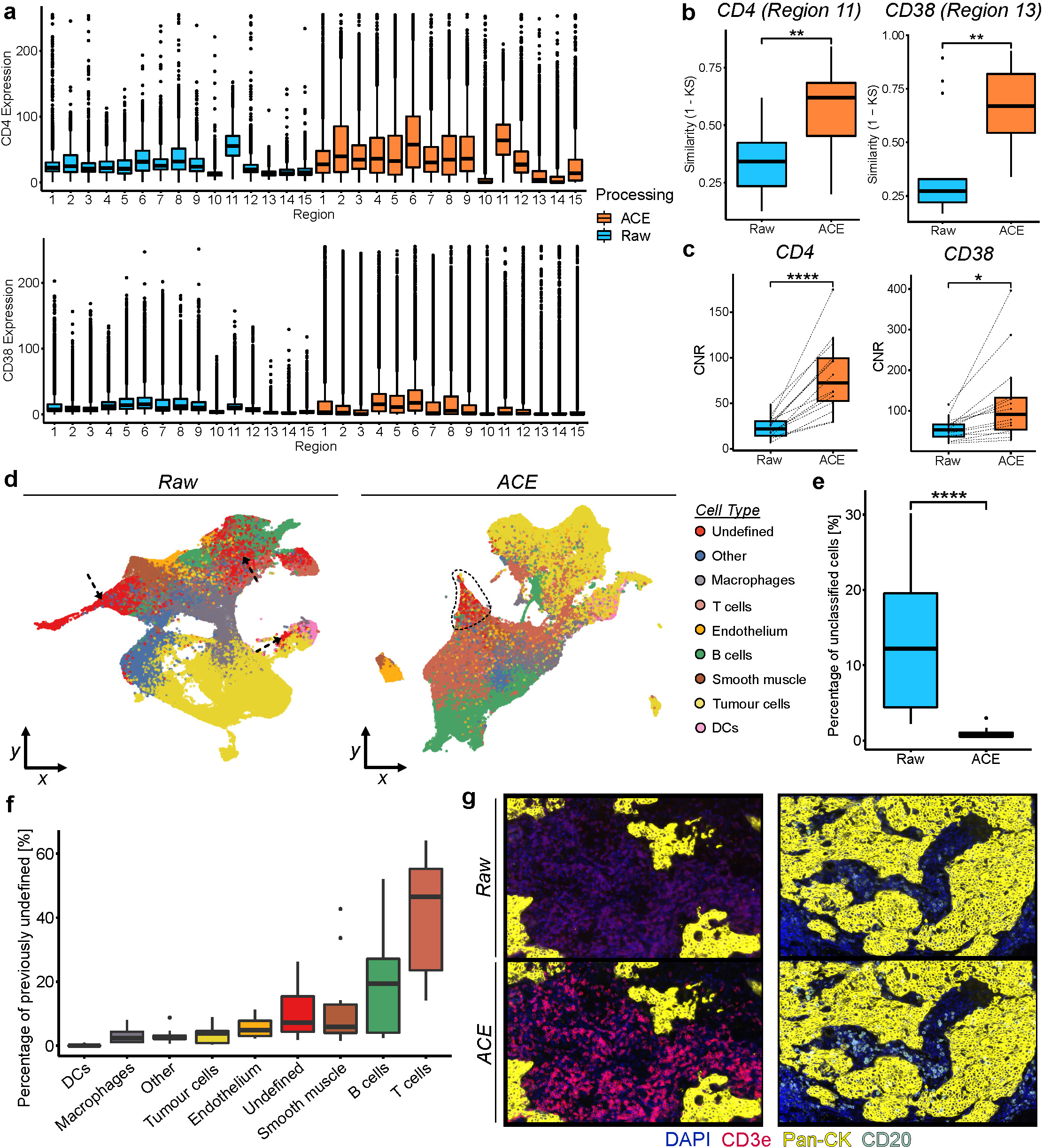
ACE normalises signal intensity across heterogeneous imaging dataset. a. Single-cell CD4 and CD38 expression per tissue core (n=15) for raw and ACE adjusted images showing a more stretched distribution on ACE data. b. Similarity of outlier cores to all other cores of CD4 (region 11) and CD38 (region 13) of raw and ACE adjusted cellular expression (n=15 for each). c. Contrast-to-noise ratio (CNR) of CD4 and CD38 of raw and ACE adjusted cellular expression (n=15 for each). d. UMAP, coloured by cell phenotype, of raw and ACE adjusted single-cell clustering and annotation showing more undefined cells in the raw data. e. Percentage of unclassified cells per core of raw and ACE adjusted single-cell clustering (n=15 for each). f. Percentage of unclassified cells per core broken up by cell type with T and B cells as the most recovered cell types (n=15 for each). g. CD3e and CD20 protein expression before and after ACE adjustment. The low expression of phenotypic markers in the raw images lead to reduced identification of these cells types. Boxplots denote the median (centre line), upper and lower quartiles (box limits), 1.5x interquartile range (whiskers) and outliers (points). The Mann-Whitney statistical test was used unless otherwise specified where p-values are denoted as; *: p <= 0.05, **: p <= 0.01, ***: p <= 0.001, ****: p <= 0.0001.

To measure the effect of signal variation on the downstream analysis and to ensure biological variation was not compromised, we clustered and phenotyped the raw and ACE adjusted NPC samples separately with PhenoGraph (Fig. 3d). Overall, ACE had fewer undefined clusters (cell types without a definitive phenotypic protein expression) presenting on the UMAP as a singular cluster. Annotation of cell phenotypes showed lineage marker protein expression associated with each cluster (Extended Data Fig. 3b). Quantifying the percentage of undefined cells across tissue cores for raw processing compared to ACE processing showed a ∼11% reduction in undefined cells (Fig. 3e). These previously undefined cells were predominately T and B cells (Fig. 3f), in keeping with CD3e and CD20 showing low expression which was corrected with ACE normalisation (Fig. 3g). To characterise the impact of these recovered cells on possible biological insights, we compared the T cell abundance of raw and ACE clustering. Significantly more T cells were identified by ACE clustering, which resulted in increased detection of CD4+ and CD8+ T cells, as well as PD1+ T cells in both CD4+ and CD8+ subsets (Extended Data Fig. 3c-e). Plotting the location of PD1+ CD8+ T cells in relation to tumour cells revealed increased T cell infiltration into the tumour bed, a critical metric for the success of immunotherapy, which would previously have been missed without ACE normalisation (Extended Data Fig. 3f).

To test the performance of ACE with other multiplex immunofluorescence imaging systems and tissues, we ran ACE on CD45 images from multiple tissue types imaged by CyCIF and COMET (Extended Data Fig. 3g-h). ACE significantly improved the CNR of images from each system comprising multiple tissue types. Furthermore, ACE outperformed other mIF based contrast enhancing algorithms across multiple channels (Extended Data Fig. 3i-j).

### LAFTA identifies known and disease specific tissue architectures

Next, we extended current spatial analyses for the identification of tissue architectures using cellular expressions, rather than cell phenotypes^6, 17^. To do this, we developed LAFTA which uses microenvironment aggregation with distance and similarity constraints to identify tissue architectures (Fig. 4a), separate to other algorithms using cellular expression^7, 21^. The similarity constraint increases weights of similar adjacent microenvironments, virtually connecting tissue components. Thus, cells get re-annotated as a function of their own expression, the distance and expression of cells in their microenvironment, as well as the similarity between anchor cells and their neighbours.

**Figure 4.**
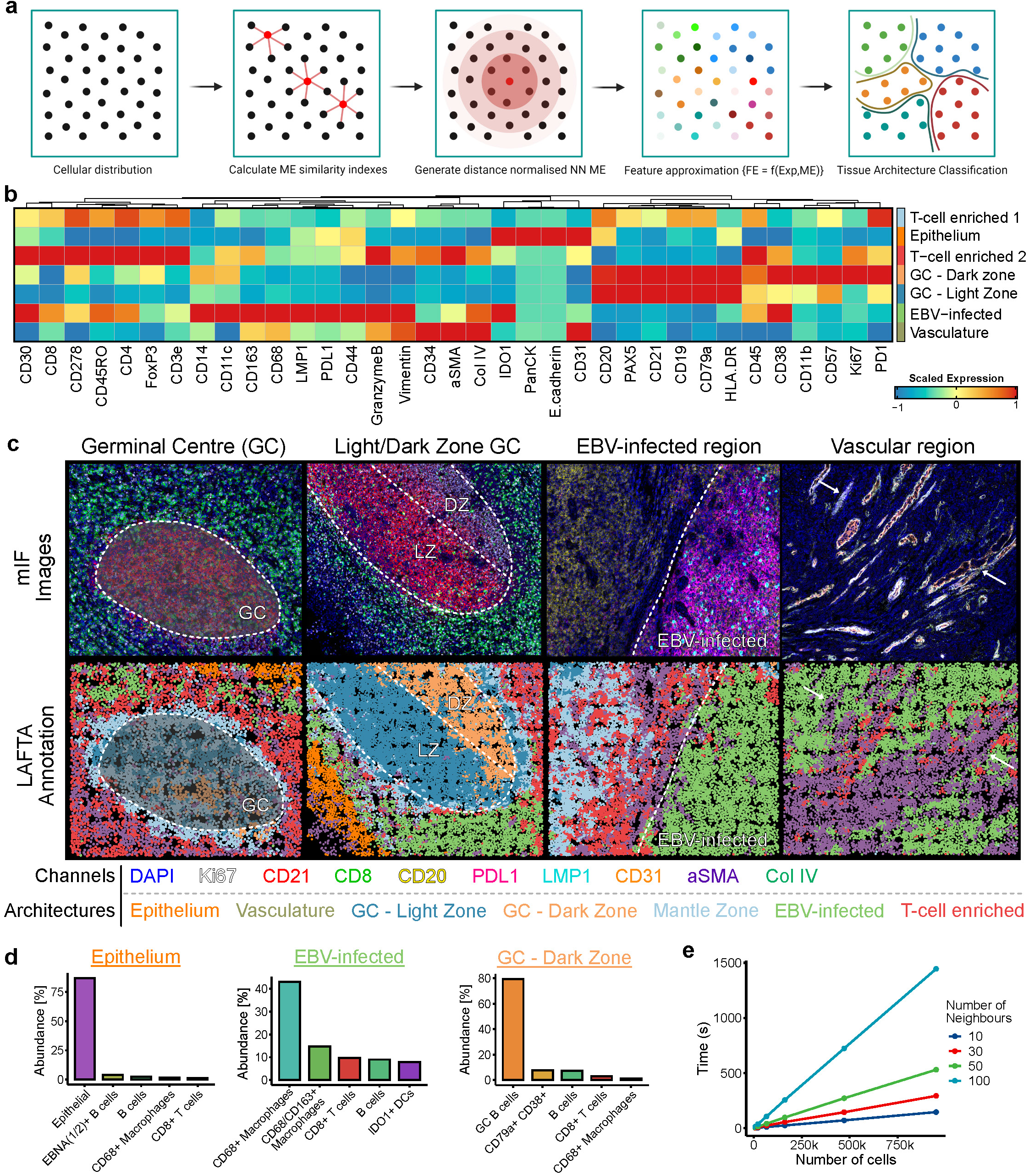
LAFTA identifies tissue architectures in infectious mononucleosis tonsil. a. Schematic illustration of LAFTA algorithm highlighting similarity and distance indexes used to develop a feature approximation map without the use of graphs. b. Enrichment heatmap of protein expression within LAFTA identifying seven IM tonsil architectures. c. Comparing mIF images and LAFTA architectures of four IM tonsil regions. d. Cell-type enrichment in epithelium, EBV-infected and dark zone germinal centre IM tonsil architectures. e. Time to calculate LAFTA architectures as a function of number of cells in the region and number of neighbour input variable.

To assess the robustness of LAFTA, we analysed regions taken from infectious mononucleosis (IM) tonsil, the symptomatic disease of primary Epstein-Barr virus (EBV) infection. We identified seven tissue structures which were manually annotated as germinal centres (GC), mantle zone, epithelium, T-cell enriched and EBV specific structures using phenotypic markers (Fig. 4b). Qualitative visualisation of LAFTA architectures showed agreement with the associated mIF images including light-zone (LZ) and dark-zone (DZ) germinal centres, without the use of specific markers for these structures (Fig. 4c). Furthermore, LAFTA captured changes due to active EBV infection including areas enriched for EBV latent membrane protein, LMP1, macrophage and cytotoxic markers and immunosuppressive checkpoint molecules (Fig. 4b & 4c). To validate LAFTA architectures, we independently clustered and annotated cells into 16 phenotypes. Quantifying the abundances of these phenotypes within each LAFTA showed the correct cell types enriched in each architecture (Fig. 4d).

Next, we tested the impact of LAFTA input variables on tissue architectures by calculating the adjusted Rand index (ARI) between manually annotated and LAFTA architectures (Extended Data Fig. 4a) ^22^. The number of cells and their relative contribution had the largest impact on ARI scores. To test LAFTA across tissue types, assay chemistries and imaging platforms, we profiled published placental samples imaged on the COMET system^23^. We identified villi, trophoblasts and fibrin-immune enriched architectures which were representative of the multiplex images and expected placental structure (Extended Data Fig. 4c,d). Next, we tested a manually annotated publicly available Imaging Mass Cytometry lung dataset across multiple cell neighbourhood/tissue architecture algorithms (Extended Figure Fig. 4e)^6, 7, 17, 24, 25^. After calculating the ARI of each region with each tool, LAFTA had significantly larger ARI that other tools tested (Extended Data Fig. 4f). The significantly increased accuracy of tissue architecture identification enables more biological insights to be extracted from traditional neighbourhood analyses. Although optimisation can be performed with LAFTA, default values showed to be identify sufficient tissue architectures across tissues, diseases and platforms (Extended Data Fig. 4b,e,g).

### The CD8 T cell microenvironment of infectious mononucleosis tonsil

To define cellular microenvironments and interactions, we present the quantification of the CD8+ T cell niche (CD8-ME, cells directly surrounding CD8+ T cells; Extended Data Fig. 5a) within tissue architectures of infectious mononucleosis (IM) tonsil defined by LAFTA (Fig. 5a), a disease characterised by robust but inadequate cytotoxic T cell responses^26^. Calculating the overall composition and expression of the CD8-ME, we found as expected, that LMP1+ B cells were enriched in the microenvironment of EBV-infected architectures (Fig. 5b & Extended Data Fig. 5b). We also saw M1-like macrophages and IDO1+ DCs enriched in this architecture. Given the heterogeneity of tissue biology, we plotted the individual CD8-MEs within EBV-infected architectures. This revealed heterogeneity with LMP1+ B cells and M1-like macrophages showing ranges of 0-30% and 0-95% at N=25, respectively (Fig. 5c). Similar differences were seen in EBNA(1/2)+ B cells, M2-like macrophages and IDO1+ DCs (Extended Data Fig. 5c). To test for common microenvironment signatures, we clustered the composition of individual microenvironments with PhenoGraph, identifying 13 clusters with unique cellular abundances (Fig. 5d,e). To visualise the differences, we plotted the enrichment of composition and expression on UMAPs, which showed a strong co-localisation of M1-like macrophages, LMP1+ B cells, PDL1 and IDO1 expression with a depletion of CD4+ T cells and other CD8+ T cells, in the EBV-infected architecture (Fig. 5f & Extended Fig. 5d,e). These microenvironments also showed high HLA-DR along with CD45RO and LAG3 expression on the anchor cells (Fig. 5g & Extended Fig. 5d). To test if neighbourhood levels impacted microenvironment architectures, we projected the composition of multiple neighbourhood levels onto UMAPs with each showing M1-like macrophage enriched regions (Extended Fig. 5 f). To quantify differences between microenvironments, we conducted differential microenvironment testing. Splitting and comparing the CD8-MEs into those enriched for, and those depleted, of LMP1+ B cells (at N=50), we calculated the effect size (ES) and the q-value (Fig. 5h). Visualising these results on a volcano plot revealed consistent pro-inflammatory processes enriched in CD8+ T cell/LMP1+ B cell microenvironments that included CD30, PDL1, IDO1 expression and M1-like macrophages, with depletion of M2-like macrophages. To examine if features are consistent across microenvironment levels (i.e. distance from the anchor cell), we re-calculated the ES for the same anchor cells with increasing cell numbers (e.g. 50, 200, 400 cells, etc.). At lower microenvironment levels (closer to the anchor cell), ES values were smaller, reflecting larger heterogeneity surrounding anchor cells or a reduced sample size. However, PD1 expression was significantly enriched in the immediate vicinity of CD8-MEs with LMP1+ B cells (ES >0.5), reducing to no effect (<0.2) at 400 cells, indicating a local effect dependent on LMP1+ B cells (Fig. 5i). Taken together, these data indicate virally driven inflammatory processes cause suppressed CD8+ T cell responses in IM tonsil, with M1-like macrophages, PDL1 and CD30 expression enriched surrounding CD8+ T cell and LMP1+ B cell microenvironments and M2-like macrophages as well as both CD4+ and CD8+ T cells depleted from these microenvironments (Fig. 5j).

**Figure 5.**
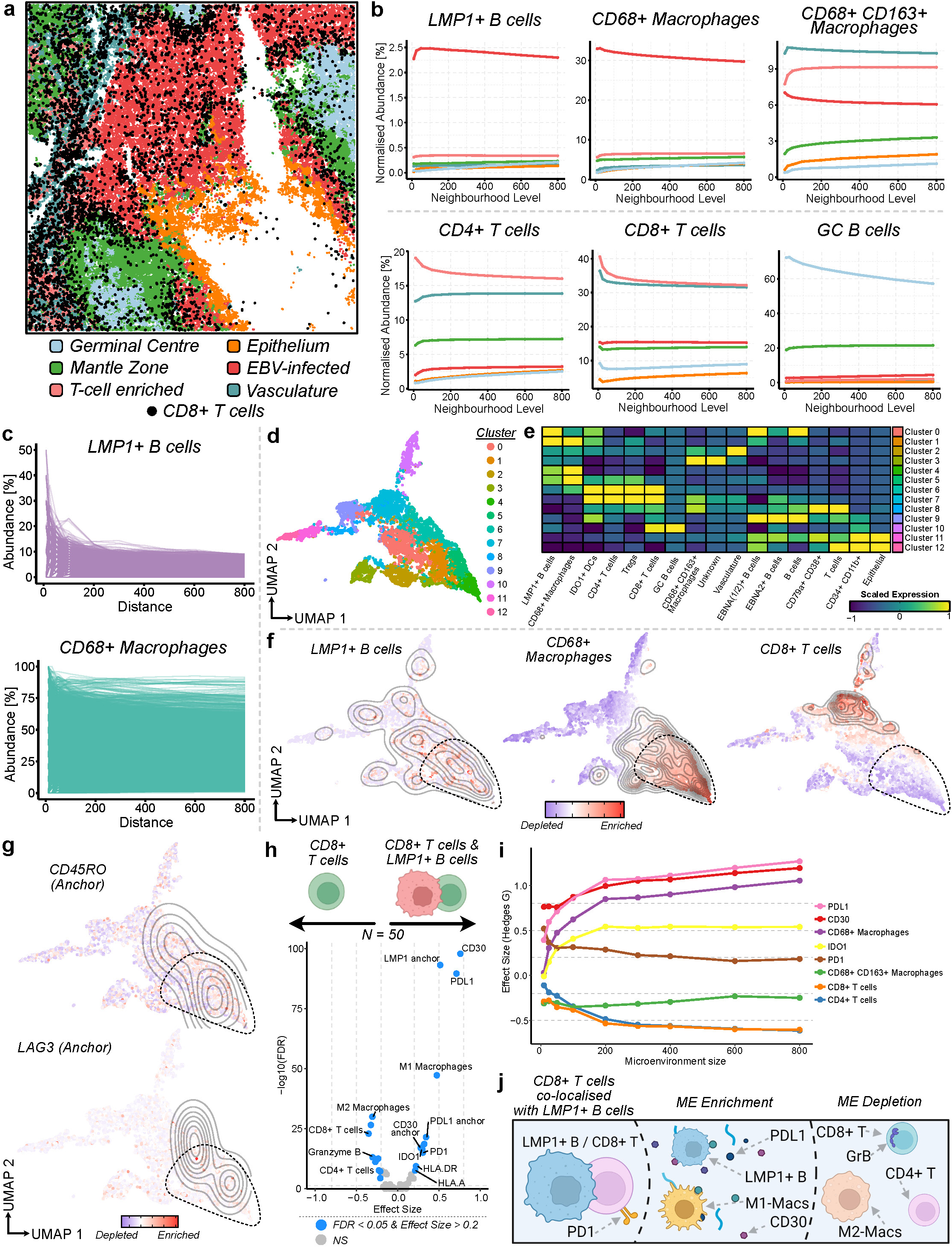
Microenvironment analysis identifies EBV-associated CD8+ T cell niches. a. Overview of CD8+ T cell locations within LAFTA architectures with reduced infiltration into EBV-infected regions. b. Microenvironment plots depicting the mean abundance (y-axis) of LMP1+ B cells, CD68+ Macrophages, CD68+ CD163+ Macrophages, CD4+ T cells, CD8+ T cells & germinal centre B cells as a function of distance (x-axis) in the immediate neighbourhood of CD8+ T cells across the different tissue architectures (colours). c. Plotting of individual LMP1+ B cell and CD68+ Macrophage compositions within CD8+ T cell microenvironments to highlight the heterogeneity within these microenvironments. d. Clustering of individual CD8+ T cell microenvironment compositions visualised on UMAP. e. Heatmap of cellular composition of each microenvironment cluster, each showing enrichment for unique cell types. f. Cellular composition of LMP1+ B cells, CD68+ Macrophages & CD8+ T cells within clustered CD8+ T cell microenvironments highlighting a link between LMP1+ B cells and CD68+ Macrophages and depletion of other CD8+ T cells in the same microenvironments. g. Anchor cell expression dependence on microenvironment composition with LMP1+ B cell enriched microenvironments having a more antigen experienced anchor CD8+ T cell. h. Differential microenvironment testing of CD8+ T cell microenvironments with and without LMP1+ B cells. i. Changes in effect size of CD8+ T cell microenvironments with and without LMP1+ B cells as a function of distance from anchor cell. j. Schematic of CD8+ T cell microenvironments in the presence of LMP1+ B cells with increased PD1 anchor cell expression and enrichment of PDL1/CD30/M1-like macrophages inflammatory microenvironment.

## DISCUSSION

A deeper understanding of spatially resolved tissues will result in improved knowledge of disease biology, which in turn will be likely to improve treatments and outcomes for patients. Currently, a lack of robust user-friendly multiplex imaging analysis pipelines is inhibiting advances in this field. Although a handful of analysis tools exist, these do not allow complete customisation of the workflow and tools, do not implement dataflow procedures, require substantial expertise to run and are optimised for idealistic experiments^20, 27, 28^ (Supp. Table 2 & 3). Hence, more robust user-friendly pre-processing tools and spatial analyses that cater for non-perfect experiments are required. We developed CellMAPS, a no-code customisable pipeline along with five new functionalities. The model-driven approach assists new users in selecting the correct input parameters while dataflow checking ensures that incompatible pipeline steps are not executed sequentially. CellMAPS is accessed through a web browser, requiring no programming experience, and can be deployed on a PC or on the cloud allowing applicability to any user. Furthermore, CellMAPS is implemented in a bespoke open architecture for biomedical image analysis, CINCO De Bio, which can be applied to other image types^29^.

As part of HIPPo, we developed three new functionalities, based on the need to develop tools that allow ease of use and the recovery of optimal numerical data from sub-optimal images, ultimately resulting in increased biological insights. To de-array TMAs for downstream processing, we developed SegArray, a user-tuneable interactive module which reduces the time associated with de-arraying of non-perfect TMAs. While de-arraying options currently exist^20^, SegArray outperformed other tools, especially on difficult to de-array TMAs (which consist of overlapping and extremely large cores). Our human-in-the-loop model further provides customisation and manual adjustment to de-array any format of TMA. To reduce technical signal variation and optimisation workload associated with running multiplex assays, we developed ACE, a multi-target regression model trained on over 9,000 images to detect upper- and lower-pixel thresholds and optimise image SNR. Although software to increase image quality in fluorescent images exists^30, 31^, ACE significantly increases SNR, reduces inter-sample expression heterogeneity, resulting in a more objective analysis pipeline. However, ACE is limited to correcting contrast limits across an entire image and hence cannot fix issues of uneven illumination or point sources of bright noise within an image (whether that be a single core of a TMA or a full-face tissue section). Yet, ACE provides an alternative to re-running costly experiments to cater for poorly performing targets. These additions to the multiplex imaging pre-processing pipeline allow users to generate optimal numerical data for downstream spatial analyses.

In MISSILe, we designed a new S4 object and developed LAFTA and microenvironment analyses tools. LAFTA, a molecular tissue architecture algorithm akin to cellular neighbourhoods^17^, uses the underlying cell expression for tissue partitioning, like UTAG and CellCharter^7, 21^, instead of cell phenotype^6, 32^. Therefore, cell annotation errors are not transferred to architecture calculation. LAFTA does not build computationally expensive graph networks or AI models, meaning it can be efficiently scaled to millions of cells^8^. A similarity constraint creates a spatially connected structure of similar cell types, with customisable settings for segregation optimisation, allowing use of k-means clustering. LAFTA can identify known and novel molecular architectures across multiple imaging systems. However, LAFTA is still restricted to only identifying tissue architectures, akin to several other tools, and does not consider acellular elements (e.g. extracellular matrix). Nonetheless, LAFTA was extended to characterise cellular microenvironments and interactions on length scales larger than direct interactions. Our microenvironment analysis uses nearest neighbour analysis, as opposed to graph-based approaches^5^, and hence is computationally efficient and scalable to millions of cells. Enrichments of cell types in microenvironments can be extended to all combinations of cell types to generate an interaction map across disease states.

In a broader context, we have shown that CellMAPS is applicable to any imaging based spatial proteomic platform, including COMET, CODEX and IMC. Operability across various infrastructures, from PC to cloud, removes reliance on specific infrastructures. In sum, CellMAPS enables broader community-wide use of a customisable multiplex image analysis suite, removing the barrier of entry to this emerging field.

## EXTENDED FIGURE LEGENDS

**Extended Figure 1.**
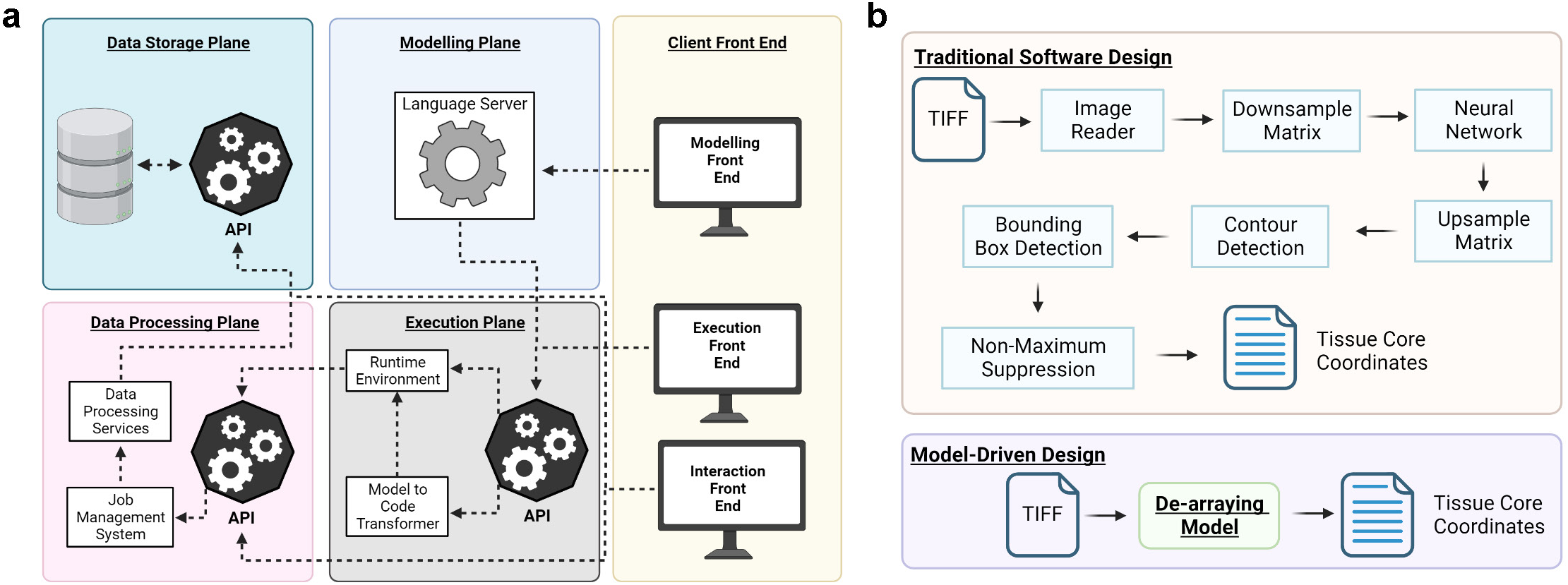
CellMAPS no-code implementation and graphical user interface. a. CellMAPS architecture implemented in bespoke CINCO de Bio architecture. b. Model-based implementation of CellMAPS. The traditional software approach is a series of functions carried out on data. The model-based approach abstracts away these functions and only concerns the user with the input and output data.

**Extended Figure 2.**
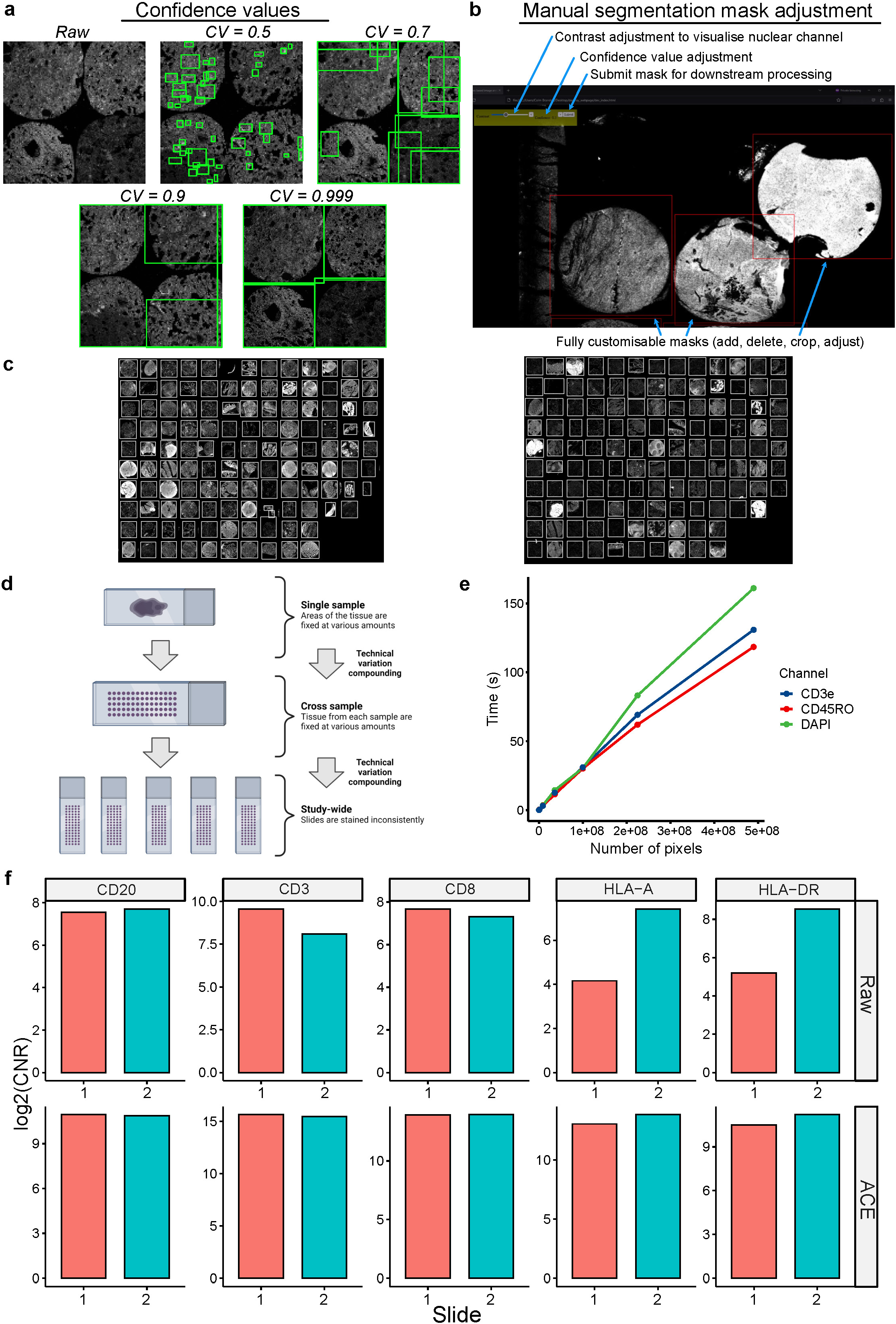
SegArray & ACE model extras. a. The impact of the confidence value variable on SegArray output of large lung cores. The increased core size and background space in the tissue (alveolar space) results in poor prediction on lower confidence values. b. Screenshot of the manual tissue de-arraying module in CellMAPS providing the user ability to adjust nuclear channel contrast, confidence value and add/delete/crop/adjust the predicted masks. c. De-arraying of two unseen TMAs with 34 additional tissue types using SegArray. d. Schematic showing the sources of signal heterogeneity and technical variations in multiplex imaging stemming from tissue fixation and inconsistent staining across batches. e. Time (in seconds) of ACE to predict thresholds as a function of number of pixels in an image. f. CNR values of raw and ACE adjusted signals for CD20, CD3, CD8, HLA-A and HLA-DR from human lymph node staining on the PhenoCylcer FUSION. Adjacent slides (Slide 1 and 2) were stained with different technical parameters. Technical variation was induced between slides by altering the antigen retrieval buffer pH, antibody mixture incubation time and time from post-staining fixation to imaging.

**Extended Figure 3.**
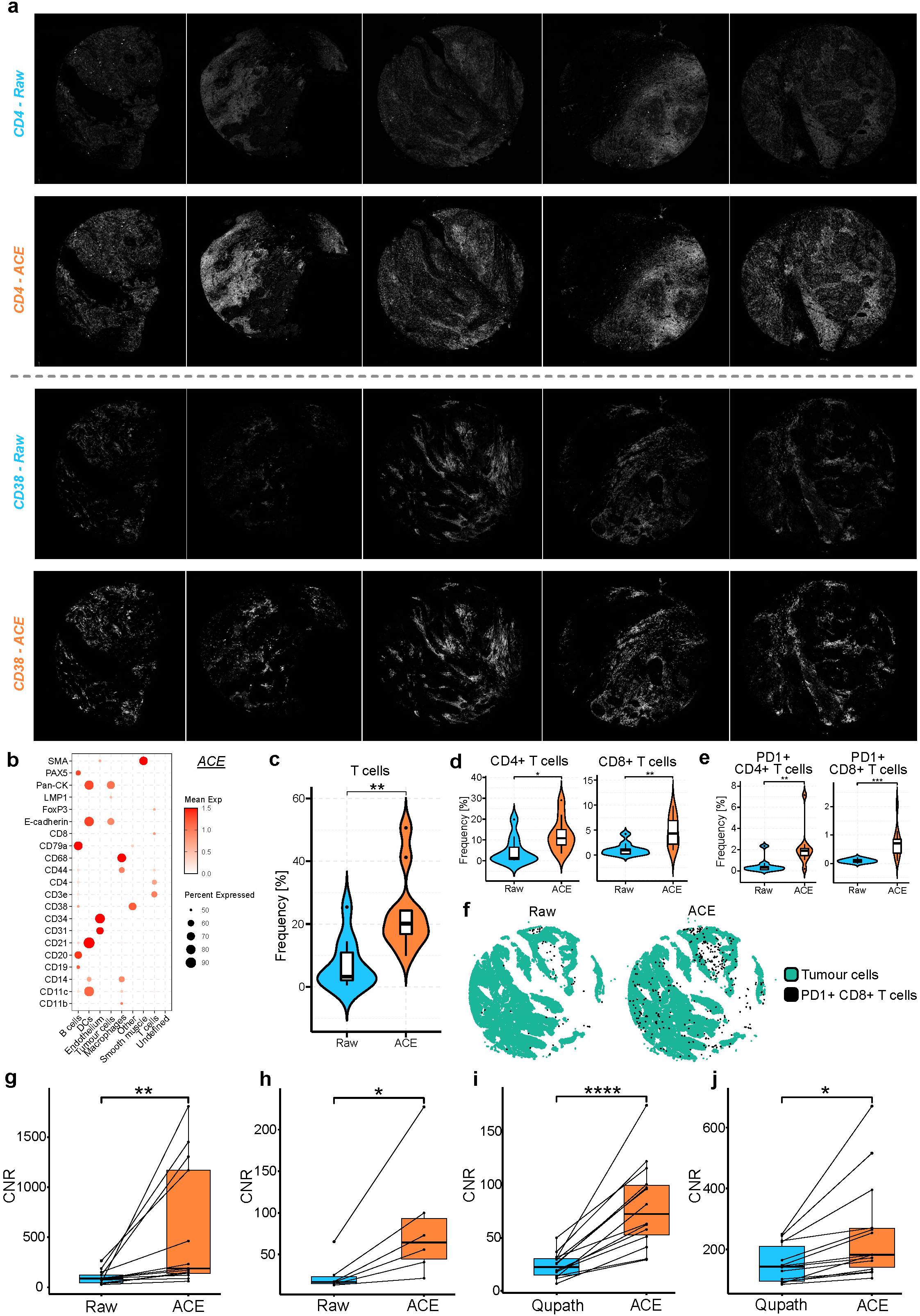
ACE benchmarking and enabled biological insights. a. Patient core heterogeneity of CD4 and CD38 before and after ACE. Visually, ACE-adjusted images show a more consistent signal across multiple cores compared to raw control images. b. Cluster protein expression used for annotation of ACE processed single-cell proteomics data. c. T cell abundance of raw and ACE annotated clustering (n=15 for each). d. CD4 and CD8 abundance of raw and ACE annotated clustering (n=15 for each). e. PD1+ CD4 and PD1+ CD8 abundance of raw and ACE annotated clustering (n=15 for each). f. Visualisation of PD1+ CD8 T cells of raw and ACE annotated clustering plotted with tumour cells (shown on the middle core from Fig. 3a). g-h. CNR of raw and ACE-adjusted CD45 images from multiple tissue types imaged by CyCIF and COMET. i-j. CNR of raw and ACE-adjusted images from multiple channels of NPC mIF imaged by PhenoCycler FUSION. Boxplots denote the median (centre line), upper and lower quartiles (box limits), 1.5x interquartile range (whiskers) and outliers (points). The Mann-Whitney statistical test was used unless otherwise specified where p-values are denoted as; *: p <= 0.05, **: p <= 0.01, ***: p <= 0.001, ****: p <= 0.0001.

**Extended Figure 4.**
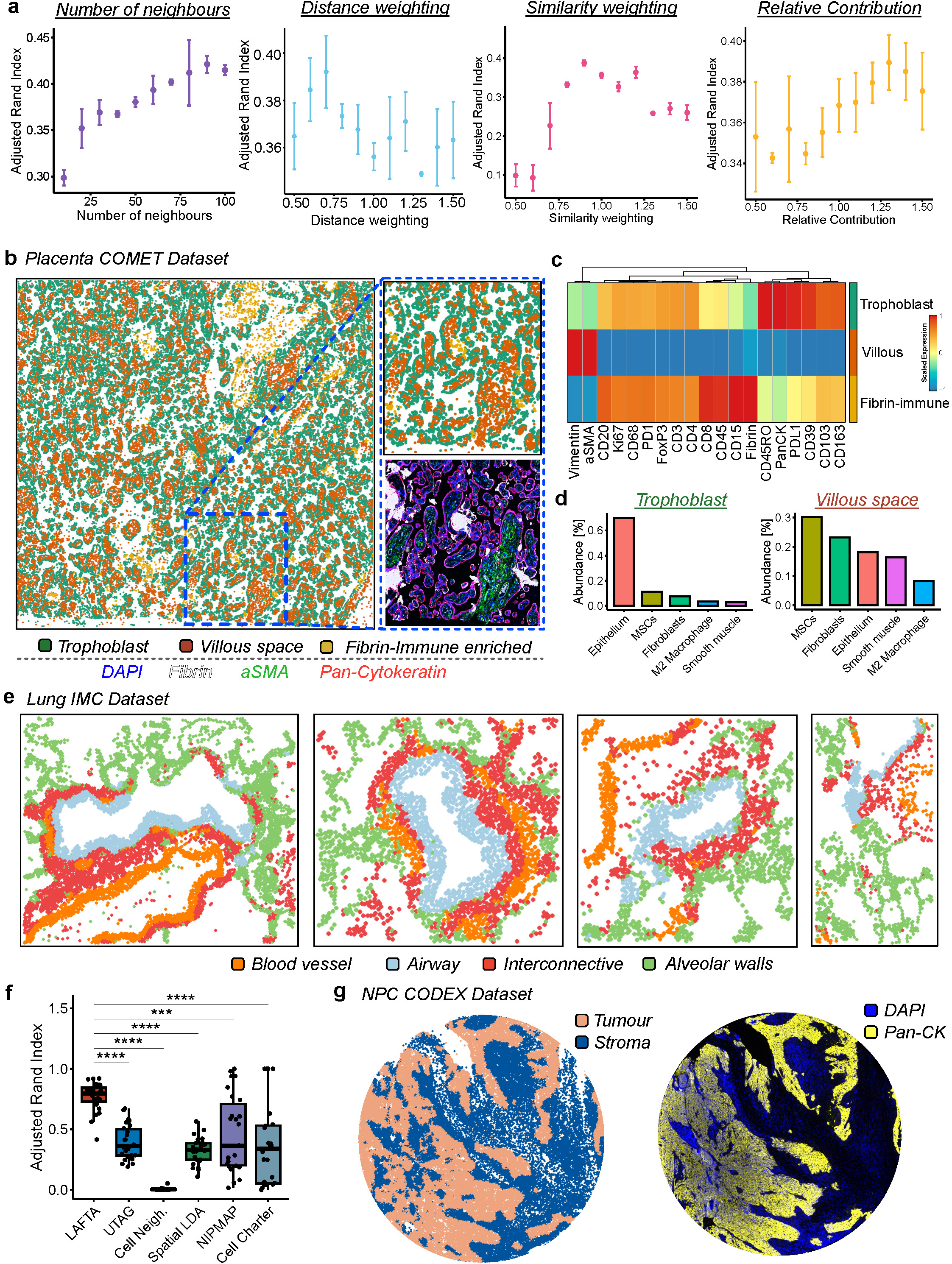
LAFTA identifies tissue architectures across platforms and panels. a. Impact of input variables (number of neighbours, distance weighting, similarity weighting & relative contribution) on LAFTA performance (error bars denote the standard deviation). b. LAFTA applied to placental tissue imaged on the Lunaphore COMET platform compared to the mIF images. c. Protein expression enrichment in identified placental tissue architectures (trophoblast, villous, fibrin-immune). d. Cell-type enrichment in placental trophoblast and villous space architectures. e. LAFTA applied to lung tissue captured by IMC identifying blood vessel, airway, interconnective and alveolar space structures. f. Adjusted Rand Index of LAFTA, UTAG, Cell Neighbourhoods, SpatialLDA, NIPMAP and CellCharter predictions from manually annotated lung IMC dataset. g. Tumour and stroma identification of NPC dataset captured on the CODEX. Boxplots denote the median (centre line), upper and lower quartiles (box limits), 1.5x interquartile range (whiskers) and outliers (points). The Mann-Whitney statistical test was used unless otherwise specified where p-values are denoted as; *: p <= 0.05, **: p <= 0.01, ***: p <= 0.001, ****: p <= 0.0001.

**Extended Figure 5.**
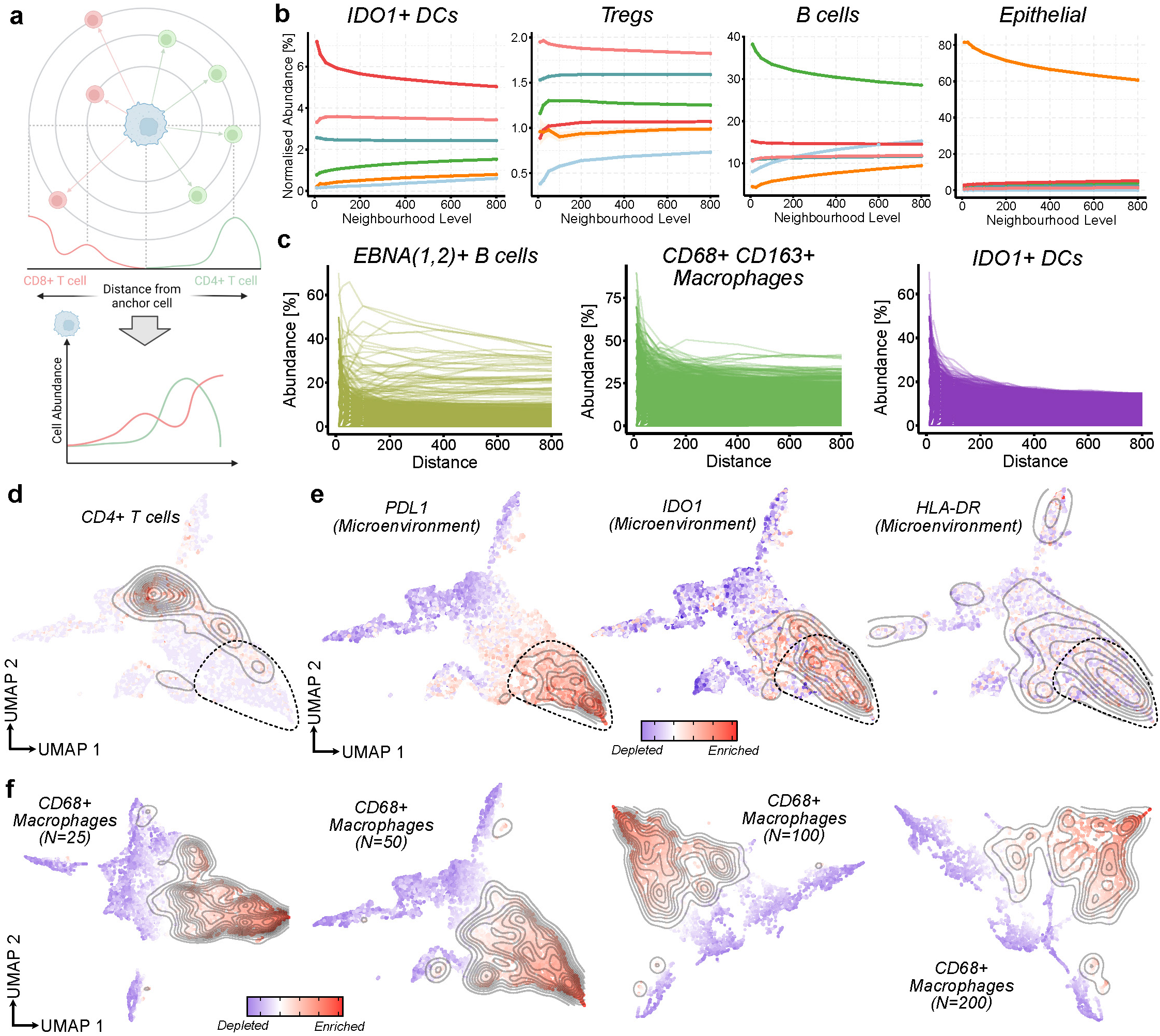
Microenvironment analysis extras. a. Schematic of nearest neighbour conversion to microenvironments. The abundance and expression of cells surrounding the anchor cells are recorded and plotted vs. distance from the anchor cell. b. Microenvironment plots depicting the mean abundance (y-axis) of IDO1+ DCs, Tregs, B cells and Epithelial cells as a function of distance (x-axis) in the immediate neighbourhood of CD8+ T cells across the different tissue architectures (colours). c. Plotting of individual EBNA1/2+ B cells, CD68+ CD163+ Macrophages and IDO1+ DCs compositions within CD8+ T cell microenvironments. d. Cellular composition CD4+ T cells within clustered CD8+ T cell microenvironments. e. Microenvironment expression dependence on microenvironment composition with LMP1+ B cell enriched microenvironments having higher expression of immunosuppressive molecules PDL1 & IDO1 as well as increased MHC Class II expression. f. Influence of neighbourhood size on microenvironment clustering showing similarity between 25 and 200 cells.

## METHODS

### CellMAPS development

#### Model-based workflow design

A core tenet of the CellMAPS system design is model-based development. Multiplex tissue image analysis integrates three domains; pathology, data analytics and artificial intelligence. As such we have developed a domain specific language and accompanying taxonomies which encode the relevant concepts from these domains in a way that is easily comprehensible by the end user. This enables the models which describe the data processing services and their inputs/outputs in a manner which is intuitive to the end user. With respect to processing services, wet laboratory end users are not typically experts in data analytics or artificial intelligence. Therefore it is favourable to user higher levels of abstraction. How each step of the process transforms the input to the output is largely irrelevant to the end user. Therefore we created models which are higher levels of abstraction of each process that perform the tasks of the underlying algorithm. Each algorithm that is integrated into CellMAPS has to conform to this higher-level model. This ensures consistency of operation for end users (e.g. if they can use one cell-segmentation model they can use them all). This same rationale is applied to all processes and algorithms which it makes to be grouped under a higher level of abstraction.

#### Integrated Modelling Environment

The Integrated Modelling Environment (IME) facilitates users to design and implement their analysis workflows in a no-code fashion. It provides a palette of components, service independent building blocks (SIBs) which can be dragged and dropped onto a canvas to design a workflow. The IME supports two types of edges between components in a workflow, control flow and data flow edges. Control flow edges define the sequences and selection logic of the workflow, i.e. the order of operation in the context of sequence logic and which path out of a set of options the workflow follows depending on some criteria in the case of selection logic. Data flow edges define how data passes between services in the workflow. The IME also actively applies static semantic and syntactic checks to user modelled workflow to ensure the current version of the modelled workflow is valid. It is important to note that the IME will prevent the user from executing a workflow unless it passes all checks, this is done to guarantee their will not be errors at runtime. The IME is implemented with the CINCO cloud framework.

#### Service Independent Building blocks

CellMAPS is composed of individual service independent building blocks (SIBs), which provide a specific functionality or service, and are designed to operate independently of all other SIBs. This enables CellMAPS to be implemented in a modular fashion, so that a set of SIBs can be used to build an arbitrary amount of unique analysis workflows with-in the constraints of each SIBs input and output requirements. Each SIB represents a standalone containerised micro-service implemented in Docker that is accessible via the CincoDeBio interface. SIBs can be developed by bioinformaticians/computer scientists and hence any third party service can be integrated into CellMAPS as long as it confirms to the process and data taxonomy. New SIBS will automatically be interoperable with the pre-existing set of SIBs.

#### Execution Environment

Once an analysis workflow is modelled in the IME it has to be transformed to executable code and subsequently executed in a runtime environment. This is handled by three components ; the execution application programming interface (API), the code generator and the execution environment. The execution API facilitates the code generation and workflow execution. Initially it handles the ingestion of workflow models from the IME and dispatches them to the code generator. The code generator takes the high-level user designed model and, using a combination of predefined templates and algorithmic techniques, generates source code in the Python/R programming. This source code represents the Python/R implementation of the user-designed workflow orchestration. The execution environment is designed to be extremely lightweight, with no data processing taking place within the execution runtime. Rather it simply orchestrates the set of standalone services (via API calls) which make up the workflow, by passing data from one service to the next, following the correct sequence and selection defined by the user. For workflow execution, the execution API acts as the interface between the data processing service and the execution runtime. The execution runtime submits the data for the current processing step to that services submission API endpoint. The execution runtime then waits to be informed by the service that it has completed processing the data and the results are then available to be retrieved.

#### Data storage

Data storage is handled using object storage. Aside from the advantages with respect to scalability and the ability to associate rich metadata which each stored data object, object storage makes data available over the network, enabling useful functionality. For example, individuals executing an analysis workflow can share final or intermediate results with a colleague who is located remotely via their web-browser. Furthermore, remote users can also execute analysis on this data. As the system does not need to handle any data directly, it enables the execution environment to be extremely lightweight. One drawback of object storage versus a mounted storage equivalent, such as block storage, is the lack of low-level access to the data. This is relevant in the context of raw experimental data which is stored in the form of a multi-page TIFF file, a typical processing step would only need to load a portion of total file (i.e. a single page) into memory. As these files are incredibly large transferring the total file over the network and subsequently loading the entire file into memory would lead to massively degraded performance. To mitigate this the system converts the raw data to an intermediate multi-file format which enables finer grained access to the data (i.e. a single page from a multi-page TIFF).

#### Implementation

CellMAPS follows the Service-Orientated Architecture (SOA) design paradigm with each component being deployed as a micro-service using containerisation. The system is deployed using Kubernetes for a cluster or minikube for single node/local deployment, both of which are available via single artefact deployment for easy install. All data processing services are implemented in the Python and R programming languages and containerised using Docker. Data storage is implemented using the S3 compatible object storage system MinIO, along with several accompanying object lambda’s written in Python. The execution and service API’s are written in Python using the FastAPI framework and conform to Open API specification v3.1 and deployed using ASGI web server uvicorn. Queuing systems are implemented using a combination of the Kubernetes Jobs API and RabbitMQ depending on the use case. Finally MongoDB is the primary database used throughout the system. The individual modules of CellMAPS, HIPPo and MISSILe can be ran independently in python or R. Full tutorials and source code are available at CellMAPS.github.io

#### SegArray

14 TMAs of varying tissue types, including lymphoid and solid malignancies, COVID-19 lung and placenta and tonsillar tissue were manually annotated using the *freehand* function in MATLAB. TMAs were down sampled five times prior to annotation using a custom MATLAB script. Varying degrees of TMA layouts were used. SegArray was trained using UNET as a starting point. To train SegArray we developed a domain specific data augmentation approach described below. Prior to training, we applied our data augmentation approach to generate 20,000 new pseudo-samples from our 13 source TMAs. Pseudo-samples were created from a blank image of arbitrary size and adding each of the background points from the real samples by randomly sampling from these points in conjunction with using some heuristics to fill up the blank side in a desired manner. Cores were then placed on the pseudo-slide background by randomly sampling a subset of the real tissue cores from all the samples. Augmentation techniques are then applied to directly to the individual objects before placing them on the pseudo-slide. We used standard computer vision techniques (resizing, rotation, contrast adjustment, etc.) as well as a new domain specific technique we developed for introducing pseudo-biological shapes to the tissue core, to simulate imperfections that are commonly seen on tissue cores. Finally the set of pseudo-cores are placed on the pseudo slide using a 2D bin packing algorithm. Along with generating the pseudo-sample the augmentation process also generates the ground truth for this sample by applying all the same operations to the mask. The images and their masks were resized to fit 1024 x 1024 pixels for the UNET model. The model was trained for 200 epochs using Intersection over Union (IoU) loss. The training and validation loss curves were monitored for overfitting and the last model checkpoint (before overfitting regime) was taken as the final number of epochs (200). When evaluating the performance of the trained model, we found that a single model would struggle to segment all cases due to heterogeneity of the data. We then explored the effect that the confidence threshold (how confident the model has to be to classify a pixel as belonging to the object class) had on the results. For examples with solid well-defined tissue cores that were highly contrasted from the background, a higher confidence threshold ∼90% was sufficient. However in samples which contained extremely low contrast and fragmented cores, a lower confidence threshold ∼50% was required. A service version of SegArray was integrated into CellMAPS which performs several predictions using multiple confidence thresholds. We then created an accompanying tool which enables the user to view these predicted ROIs overlayed on the original image in their browser, cycle through them by confidence threshold, select the best predicted set and make any necessary adjustments to the ROIs. Tissues were then cropped by the smallest bounding box including the whole core. If there is no bounding box small enough to contain only a single core (i.e. another core lies somewhere outside the perimeter of the first core but within the area of the box centred on the first core), that portion of the second core will be included in the crop.

#### Data Augmentation

To address the scarcity of annotated TMA images and enhance robustness in dearraying models, we developed a novel data augmentation pipeline. This method synthesises TMA slides from real source data while preserving pixel-level ground truth for use in model training. The method begins with organic shape synthesis, which generates natural-looking closed contours by distributing control points uniformly on a randomly sized circular base, then applying controlled perturbations to introduce asymmetry. These modified points undergo cubic spline interpolation with periodic boundary conditions, creating smooth, continuously differentiable closed paths that are rendered as filled silhouettes. Geometric transformations (e.g. rotation, shearing, etc.) further enhance realism to mimic Tissue Microarray (TMA) imperfections like core fragmentation or voids. Next, for synthetic slide generation, pixel intensity distributions from source TMA images are sampled to create backgrounds, optionally blended from multiple sources for heterogeneity. Each tissue core undergoes randomised transformations—including contrast adjustments, non-uniform scaling, rotations, and insertion of the organic-shape imperfections described above. Then a 2D bin packing algorithm positions the set of cores without overlap, preserving natural spatial relationships through a greedy approach (with heuristic methods planned for future exploration). The final pseudo-synthetic TMA is then created by integrating the transformed cores with the synthesised background. Throughout the pipeline, identical transformation sequences are applied to both tissue cores and their corresponding binary masks, ensuring pixel-perfect ground truth correspondence across all augmentation stages.

#### ACE

ACE was trained using over 9,000 image tiles from 8 proteins encapsulating immune cells, epithelial cells, endothelial cells and functional markers, with signal intensities ranging from high to low across 3 tissue types. Tiles were created using the *patchify* library in python. Tiles were imported into QuPath and adjusted manually by experienced mIF users to the optimal qualitative signal-to-noise ratio. The lower and upper threshold pixel values were recorded. The image tiles were subsequently adjusted using a custom adjustment function in python and saved to file as new tiff images. The adjusted images were used as the ground truth for the ACE model. ACE is a multi-target regression model with the raw normalised histograms (0-1) as input and the adjusted images normalised histograms as ground truth. ACE predicts upper and lower adjusted image thresholds and then the images are subsequently altered to these values. Normalising the histogram allows the prediction to be agnostic to the size of the input image and hence the model can be trained on tiles (sub-sections of a larger image) and used for inference on full sized images with consistent results. Several regression algorithms (Ridge, Linear, Bayesian, Multi-Layer Perceptron, Gradient Boosting, Random Forest, Histogram Gradient Boosting and Support Vector Regression) were evaluated using a Grid Search across a set of best-practice hyper-parameters for each algorithm and 5-fold cross validation performed on the training set to give a clear indication of which algorithm was performing optimally. Random Forest Regression was the best scoring model with a mean R^2^ (coefficient of determination) value of 0.82 on the cross-validation sets. The model was then trained on the 5-folds and evaluated on a separate hold-out test set (which contained patches of protein expressions not in the training set) achieving an R^2^ value of 0.76. ACE was implemented using Scikit-learn. Other methods were also explored to implement a contrast equalised, including pixel-to-pixel generative adversarial networks (insufficient data) and a genetic algorithm with a neural network-based fitness approximation with varying degrees of success (inconsistent results). However, multi-target regression provided the best results. Although ACE is designed explicitly for use on images with 8-bit colour resolution, 16-bit colour images are down-sampled prior to inference.

#### ACE Benchmarking and Implementation

To benchmark ACE predictions on tiles, 320 unseen image tiles across eight proteins and two tissues were manually adjusted, and the lower and upper thresholds recorded. ACE predictions were then made on these tiles. The cosine similarity between manual thresholds and ACE predictions (lower and upper) was then calculated for various proteins. The architecture of ACE allowed a seamless transition from image tiles to full core/face tissues. We qualitatively validated that the large images were being adjusted correctly.

#### Cell Segmentation

CellMAPS includes native options to segment cells using CellSeg, DeepCell or CellPose 2.0 ^11–13^. For CellSeg, a nucleus-based segmentation that expands the membrane by a chosen number of pixels, the nuclear channel will be taken as the first stack in the tiff image. DeepCell and CellPose can be used as nucleus-based or membrane-based segmentation tools. In the case of membrane-focused segmentation, a membrane channel must be selected. CellMAPS allows the user to select a stained membrane marker (either a dedicated option, e.g., Sodium ATPase, or a lineage marker that is present in most cells, e.g., CD45 in lymph node). However, CellMAPS also allows generation of a *pseudo-membrane* marker by combining several markers together from various protein channels which stain different cell types in heterogeneous tissue (e.g., Pan-cytokeratin, CD45, CD31, etc. in breast cancer). DeepCell and CellPose make predictions on this pseudo-membrane and provide a discretised segmentation mask which marks the pixels for each cell. They also make predictions on the nuclear channels which CellMAPS uses as the first stack in the tiff image. As membrane-based segmentation is more accurate but less robust than nuclear based segmentation, some cells will not be segmented adequately. For these cells, CellMAPS calculates which membrane masks contain multiple nuclei masks (which usually indicates doublets) and deletes the associated membrane mask. These nuclei are then expanded as is the case in nuclear based segmentation. This combines the accuracy of membrane-based segmentation with the robustness of nuclear based segmentation.

#### Xtractit

To extract expression and morphological features from the multiplex images, *regionprops* (scikit-image) is used to define the pixel coordinates and morphological features (e.g., size, eccentricity, etc.) of both the membrane and nuclear mask from each single-cell. The pixel coordinates are then used to calculate either the average expression of the raw or log2 normalised pixel intensities. For nuclear channels, the user defines the nuclear markers in the panel (e.g., FoxP3, Ki67, etc.) and the extraction is completed from the nuclear masks rather than membrane masks. This ensures the nuclear expressions are only normalised by the nuclear areas. The extracted data is formatted as a csv/fcs file where each row is the information associated with each cell and the columns are the morphological/expression data. The csv/fcs files can further be applied to REDSEA, a spatial spillover compensation algorithm, to further clean the numerical data^14^.

#### MISSILe

Multiplex Immunofluorescence Spatial Interaction Library (MISSILe) has been developed for spatial analysis of multiplex image data. It has been integrated into CellMAPS but can also be used as a standalone R package. MISSILe is centred around a S4 class MISSILe object with slots for counts, spatial information, metadata, neighbour interactions, neighbourhoods/architectures, microenvironments and dimensionality reductions. Functions within MISSILe are designed to update these slots with the corresponding analysis. Within MISSILe, cell quality can be controlled by filtering cells by various morphological and expression thresholds. Phenotyping can be done by unbiased clustering using FastPG, allowing analysis of over 2M cells^33^. However, CELESTA can also be used to phenotype cells with a user generated input matrix of expected cell types and expression profiles ^16^. Cell-to-cell interactions and cellular neighbourhoods can be calculated with several metrics and plotted using heatmaps or boxplots. MISSILe includes multiple visualisation and plotting functionalities including cellular abundances, spatial expression, spatial phenotypes, and others.

#### LAFTA

LAFTA is used to define structures within tissue with similar molecular characteristics. Raw cellular protein expression, extracted from Xtractit and imported as a *csv* file into MISSILe, is used as input to LAFTA. The N nearest neighbours (termed microenvironment) are calculated for every cell (anchor cell) using nn2 (from the RANN R package). The similarity of the microenvironment of each cell is calculated (using the Euclidean distance) against all its neighbour cells, generating a connected map of the tissue without the need for computationally expensive graphs. Cells within each microenvironment are then normalised by their distance from the anchor cell ensuring cells far away do not impact the LAFTA calculation as much as those close. The similarity and distance calculations generate a feature approximation score of each cell which is then clustered into bins that represent tissue architectures. For speed, we implemented the clustering in K-means clustering as multiple millions of cells may have to be profiled. Several values of K can be screened to identify the optimal number and clusters can be combined for increased accuracy. Examining run time, LAFTA calculated one million cells in four minutes with 30 cells per microenvironment (Fig. 4e). LAFTA results can be visualised on a heatmap of clusters against protein expression and the spatial point patterns of each tissue coloured by LAFTA architecture. Furthermore, an essential validation step to assist in the interpretation of LAFTA architectures is plotting the enrichment of annotated cell types within each architecture.

#### LAFTA Manual Annotations

Given the difficulty of annotating IM tonsil due to its altered structure^22^, we manually annotated a singular region with minimal EBV-infected B cells. Regions were drawn by a consultant pathologist (M.R.P.) using QuPath into epithelial, mantle, light-zone germinal centre and dark-zone germinal centre and interfollicular regions. Masks were generated for each region and imported into MISSILe. Cells in this region were subsequently assigned to one of the annotated regions. The adjusted Rand Index was calculated between annotated and LAFTA calculated architectures for varying LAFTA input variable values.

#### Microenvironment analysis

The microenvironment analysis is an extension of LAFTA to characterise the microenvironments of anchor cells. Using nn2, the cell abundance and expression of the microenvironment of the anchor cells (e.g., tumour cells) are calculated at increasing distances (e.g., the default is 25, 50, 100, 200, 400 & 800 cells). These calculations are then plotted as line graphs of abundance/expression as a function of distance from the anchor cell using the mean and standard error. Given the heterogeneity of these microenvironments, our analysis allows the user to select a microenvironment distance level and cluster the microenvironments based on cellular abundances. A UMAP and associated heatmap is then generated for each anchor cell (based on microenvironment rather than expression) showing the various unique clusters. These UMAPS can be coloured by the abundance of cells or expression of a protein in the microenvironment or the expression of the anchor cell to enable cellular enrichments and interactions to be visualised. To quantify these enrichments, we developed differential microenvironment testing to compare clusters or high/low microenvironments. The effect size (Hedge’s G) and q-value (Wilcoxon test followed by Bonferroni adjustment) of the difference between cell abundances and expression in compared microenvironments is calculated and plotted on a volcano plot. Interesting findings can be carried forward and tested as a function of distance. This method can be applied to compare clusters, regions that are high or low for cell types or cellular expression, or co-localised cells (e.g., CD8+ T cells in the presence of LMP1+ B cells or not). Furthermore, differential microenvironment testing can be applied tissue-wide to quantify enrichment of interactions between cell types on larger length scale than directly touching.

### Experimental Methods

#### Tissue acquisition and processing

A FFPE Nasopharyngeal Carcinoma (NPC) TMA, comprising 15 cores and 10 patients, and FFPE Infectious Mononucleosis (IM) tonsil tissue were obtained from the University of Erlangen, Germany. Anonymised FFPE tissue was transferred to the University of Birmingham for further laboratory experiments under the LoST-SoCC study (IRAS 193937) following approval by the ethical research ethics committee (19/NE/0336). For PhenoCycler FUSION mIF staining and imaging, blocks were cut onto positively charged frost-free slides within a 35 x 16mm window. Slides were then transferred to Akoya Biosciences, Menlo Park, California, USA.

#### PhenoCycler FUSION Staining & Imaging

A 42-plex PhenoCycler FUSION panel was designed, validated and deployed at Akoya Biosciences, Menlo Park, California, USA. The panels, clones and barcodes used are available in supplementary table 1. Glass slides were prepared, stained, and fixed as per the PhenoCycler protocol. Briefly, slides were deparaffinised in xylene and re-hydrated in decreasing concentrations of ethanol (100%, 90%, 70% & 50%). Heat Induced Epitope Retrieval (HIER) was performed in a pressure cooker at 110°C for 18 minutes in Tris-EDTA buffer (pH = 9.0). The tissue was then incubated in PhenoCycler Staining Buffer (Akoya Biosciences) for 30 minutes to block non-specific binding of antibodies. Subsequently, the tissue was incubated in a cocktail of the conjugated antibodies overnight at 4°C, then fixed in 4% paraformaldehyde for 10 minutes, 100% methanol for 5 minutes, Fixative Reagent (Akoya Biosciences) for 20 minutes and stored until imaging. Akoya Reporters were added to the corresponding well of a 96-well plate in preparation for imaging, based on the cycle design of the experiment. Autofluorescence subtraction, stitching and compression were completed in the FUSION software resulting in .qptiff files in preparation for processing and analysis.

### Analysis Methods

#### Pre-processing of NPC and IM tonsil tissues

NPC and IM tonsil tissues were processed through the HIPPo portion of CellMAPS to generate single-cell protein expression data. SegArray was used to de-array the NPC TMA. ACE normalisation was used on a per core/tissue basis for both diseases. However, for comparative purposes, the NPC TMA was also pre-processed fully with no ACE normalisation. CellSeg segmentation was used for both NPC and IM tonsil images. Single-cell protein expression *csv* files were imported into MISSILe.

#### ACE metrics

To test the core-level normalisation of ACE for the single-cell protein expression levels, we selected region 11 for CD4 and region 13 for CD38 as outliers based on boxplot values and qualitative image scoring. Taking the single-cell expression data as distributions, outlier regions were then compared to all other regions for both ACE and raw processing using the Kolmogorov–Smirnov (KS) divergence test, and subsequently subtracting the KS scores from 1 to get similarity. Contrast-to-noise ratio (CNR) was calculated using (Sig_mean_ – Background_mean_) / Background_SD_ for CD4 and CD38 and the values obtained were compared between ACE processed and and raw images.

#### Statistics

The Wilcoxon test was used throughout the manuscript unless otherwise specified. p-values were adjusted using the Bonferroni correction where required. Hedges G was used to calculate effect size of each feature between microenvironments.

## DATA AVAILABILITY

All data generated for this study will be made available upon publication. A multiplexed image to test CellMAPS with can be downloaded here https://zenodo.org/records/13836803.

## CODE AVAILABILITY

The software and source code used to generate and run CellMAPS are available at https://www.github.com/EannaFennell/CellMAPS.

## ACKNOWLEDGEMENTS

The authors would like to thank Akoya Biosciences for imaging the NPC and IM tonsil samples. É.F. is funded by an Irish Research Council Postdoctoral Fellowship (GOIPD/2022/97) and HORIZON-TMA-MSCA-PF-GF (no. 101110227). Illustrations were created with BioRender.com.

## AUTHOR CONTRIBUTIONS

É.F. & C.B. conceived and designed the study;

M.R.P & G.N obtained ethical approval for and acquired the tissue samples;

N.N & A.H. stained tissue samples and acquired multiplex imaging data;

É.F. developed the original algorithms and pipeline in python and R;

É.F. developed LAFTA and microenvironment analysis;

É.F., A.R. & C.L. annotated images for ACE training;

C.B., A.S. & É.F. developed SegArray & ACE;

C.B., S.R. & A.S generated the SIBs for HIPPo and MISSILe;

C.B. & S.B. generated the no-code execution environment;

É.F. & C.B. prepared all the figures;

É.F., C.B., T.M. & P.G.M. wrote the manuscript;

G.S.T., O.B., T.M. & P.G.M. supervised the work.

All the authors contributed to the final version of the manuscript.

